# Predicting phenotype to mechanotype relationships in cells based on intracellular signaling network

**DOI:** 10.1101/2023.03.24.534160

**Authors:** Esra T. Karabay, Amy Turnlund, Jessica Grear, Stephanie I. Fraley, Parag Katira

## Abstract

Cells originating from the same tissue can respond differently to external signals depending on the genotypic and phenotypic state of the cell and its local environment. We have developed a semi-quantitative-computational model to analyze the intracellular signaling network and its outcome in the presence of multiple external signals including growth factors, hormones, and extracellular matrix. We use this model to analyze the cell’s mechanical response to external stimuli and identify the key internal elements of the network that drive specific outcomes within the response space. The model is built upon the Boolean approach to network modeling, where the state of any given node is determined using the state of the connecting nodes and Boolean logic. This allows us to analyze the network behavior without the need to estimate all the various interaction rates between different cellular components. However, such an approach is limited in its ability to predict network dynamics and temporal evolution of the cell state. So, we introduce modularity in the model and incorporate dynamical aspects, mass-action kinetics, and chemo-mechanical effects on only certain transition rates within specific modules as required, creating a Boolean-Hybrid-Modular (BoHyM) signal transduction model. We present this model as a comprehensive, cell-type agnostic, user-modifiable tool to investigate how extra-and intra-cellular signaling can regulate cellular cytoskeletal components and consequently influence cell-substrate interactions, force generation, and migration. Using this tool, we show how slight changes in signaling network architectures due to phenotypic changes can alter cellular response to stress hormone signaling in an environment-dependent manner. The tool also allows isolating effector proteins driving specific cellular mechanical responses. Ultimately, we show the utility of the tool in analyzing transient chemo-mechanical dynamics of cells in response to time-varying chemical stimuli.

## INTRODUCTION

In this work, we present a deployable computational tool to connect and analyze extra-cellular signaling, cellular phenotype, and their combined influence on determining cellular mechanotype. Cellular mechanotype, as defined by cellular mechanical properties such as stiffness, adhesion strength, and contractility, among others, drives cell-extracellular matrix interactions^1–4^. These interactions determine the outcome of several key biological processes such as wound healing, metastasis, and inflammation^5–7^. The cellular mechanical properties are a function of the organization and activity of key cytoskeletal elements such as actin, microtubules, intermediate filaments, associated motor proteins, and kinases, and their coupling with cell membrane proteins such as integrins and cadherins^8–13^. The organization and activity of these cytoskeletal and membrane proteins is in turn governed by complex biochemical signaling pathways within the cell that integrate various cell intrinsic and extrinsic signals^14–17^. Our understanding of how cellular protein expression levels, the availability of various metabolites and the presence of external stimuli influences cellular cytoskeletal organization and the cell mechanotype is limited.

Mapping the biochemical reaction pathways associated with the regulation of cellular cytoskeleton can provide an estimate of changes in the availability, activity, and organization of the various cytoskeletal elements. However, such mapping is complicated due to the presence of several signaling cross-talks and compensatory mechanisms across the signaling elements ^18,19^. A deep literature analysis of the various biochemical cues that influence cytoskeletal elements reveals a massive interactive network of signals with several complex interactions that need to be considered to obtain a complete picture of cytoskeletal regulation. Figure 1A and Figure S1 (elaborated network) depict a comprehensive and intertwined network with cytoskeleton entities. Incorporation of these detailed and complex interactions across various cytosolic signaling elements and their links to cytoskeletal organization is critical to explain several unique cellular mechanical responses seen in biology. For instance, stress hormones commonly attenuate breast cancer cell migration ^20^ while increasing migration in highly metastatic breast cancer cell phenotypes ^21,22^. This phenomenon requires further research on how stress hormones induce opposite mechanical responses in cellular subtypes. Additionally, there is limited understanding of how the response to stress hormone signaling may depend on the presence of other competing signals, for example those coming from the extra-cellular matrix or neighboring cells via paracrine or juxtracrine signaling. As another example, kinase inhibitors that impact cell contractility and consequently cell migration are proven to be effective in clinical models of tumor growth^23^. However, due to the presence of competing kinases and their altered activities within different environments, targeted therapeutic inhibitors cannot entirely block these migratory pathways, which later leads to drug resistance, survival and dispersal of cancer cells ^24–26^. Understanding how these inhibitor targets coordinate the cytoskeleton and lead to a certain cellular response can provide a significant increase in the efficiency of drug administrations. Computational models can enable analyzing these complex chemo-mechanical signaling pathways efficiently ^27–30^. However, most models aimed at connecting biochemical reactions to cellular cytoskeletal organization and mechanics, so far, focus on only a limited number of signaling pathways and biochemical interactions.

**Figure 1.**
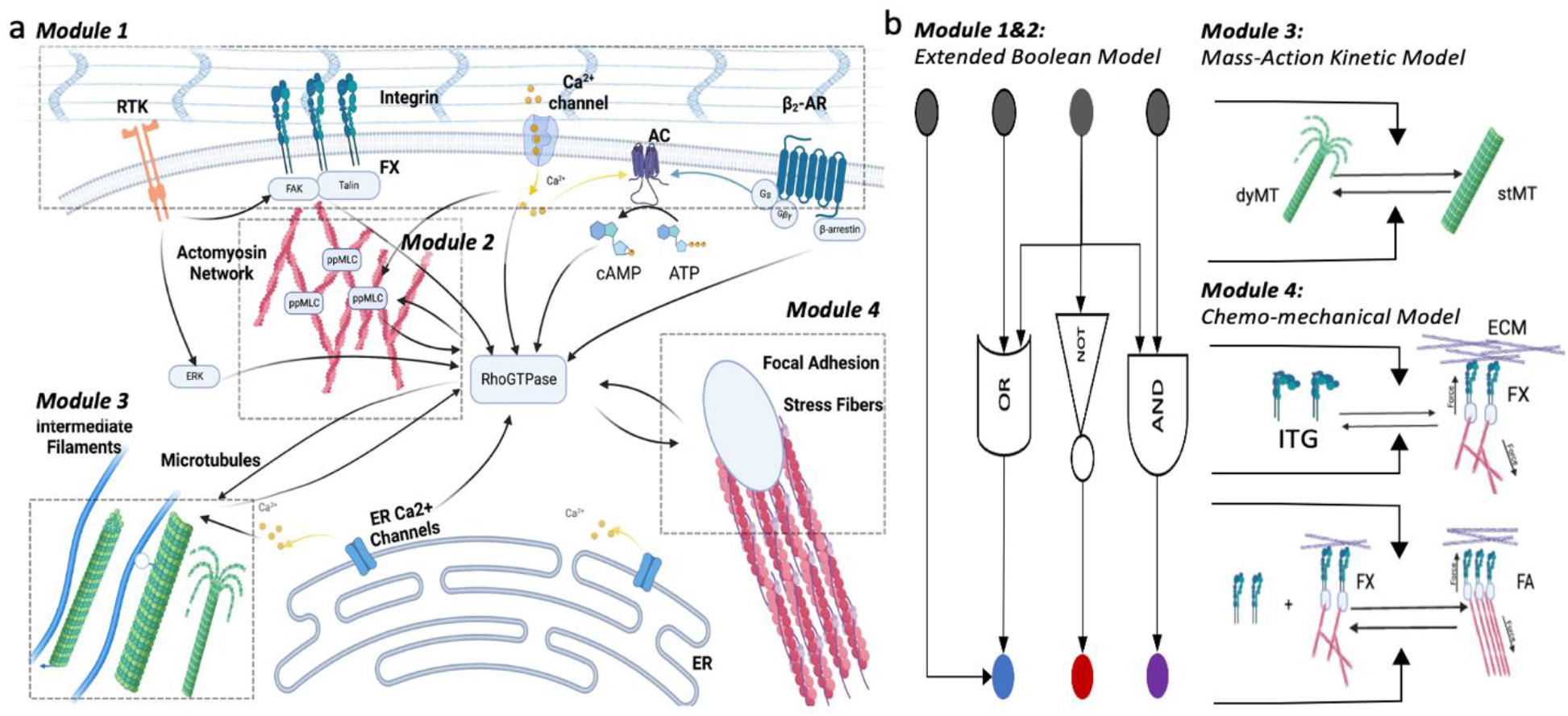
Signaling between cytoskeleton modules and plasma membrane receptor and its Boolean Hybrid Modular Model (BoHyM). (a) Network involving receptor tyrosine kinases (RTKs), β_2_-adrenergic receptor (β_2_AR), Integrin (ITG), Calcium channels and cytoskeleton elements. Actual networks consist of 120 network components, and all detailed networks can be found in the supplementary. (b) Module 1&2: Signaling networks computed with extended Boolean model which does not require kinetic parameters. Gray nodes are plasma membrane receptors and blue, red and purple nodes represent cytoskeletal-ECM anchoring moieties. Module 3: Mass-action kinetics used to compute microtubule signaling which requires mass conservation between microtubule dynamic states. Module 4: Chemo-mechanical model to capture focal adhesion formation and maturation in response to mechanical force generation by the cell and from the environment. Modules 3 & 4 receive updates from Module 1 & 2 with the extended Boolean model, in return, feedback the associated local dynamics to modify specific Model 1 & 2 node activation states.

Signaling networks can be modeled in several ways. The most common approach is writing ordinary differential equations that track the state of each element in the network over time. This approach however requires a detailed knowledge of the rates at which different interactions in the network let the state of interacting elements evolve. For large networks with several tens or hundreds of elements and many more interactions, knowing all these rates is impossible. Alternatively, Boolean models that allow only two states for each element - 0 (inactive) or 1 (active) can simplify large-networks into simple logical outcomes. Such Boolean models have been successfully deployed to describe a wide range of processes such as aging ^31^, tumor promoter signaling ^32^ and T-cells dynamics ^33^. Boolean network models can be expanded by using ODEs which enable the observation of temporal changes upon perturbations ^34,35^ and give flexibility to remodel the network associated with specific cell types by controlling signal growth and decay rates. Although Boolean models are convenient for analyzing the dynamical states of modules in large networks, there are two main challenges to achieve realistic results in biological network simulations using this approach. First, in a Boolean network, every node changes the status of the succeeding node at the same time, whereas in realistic signaling system it happens asynchronously depending on presented reaction time and catalysis. Secondly, nodes when depicted as switches can be on/off anytime. However, if nodes represent concentration, “mass conservation” has a limitation on binary logic. For example, consider a simple feedback loop, A → B ⊸ A. We assume that initial values of A-B are 1-0, which then follow the 10-11-01-00-10 pattern and continue with the same repetitive states. This is an issue unless there is limitless source for A, otherwise B inhibition cannot switch on “A” again since there would be no A. Another example of Boolean limitation is where A is directed to multiple nodes, A → B, A → C, A → D. In a conventional Boolean network approach, the signal from “A” does not get divided across the succeeding nodes states but rather affects them all. This is again problematic from a mass conversation perspective if usage of A along one pathway limits its availability for another.

As mentioned before, Boolean models can be extended with differential equations. Additionally, large Boolean networks can be coupled with other modeling approaches such as Flux Balance Analysis^36^ where specific signaling compartments of the large network are analyzed using a different, more detailed approach and the outcomes of these specific modules are integrated back into the large Boolean Network model. Extended Boolean models alleviate simulation of biological network dynamics and are helpful to overcome some of the challenges in binary world. In order to capture cytoskeleton signaling, we present Boolean Hybrid Modular (BoHyM) model involving extended Boolean function, chemo-mechanical and mass-action kinetics (Figure 1B). Using this model, we can capture how (i) competing biochemical signals can lead to diverse mechanotypes in different environments and altered intracellular signaling network architectures, (ii) breakdown how individual pathways may contribute to specific cytoskeletal changes and compensate or compete, and (iii) how transient signals drive the oscillatory mechanical response. Each part reveals unique usage of the BoHyM model to link the biochemical signals to cell mechanical elements. Overall, we provide a modifiable and easily deployable BoHyM modeling resource for researchers to analyze their own biochemical-mechanical signaling cross-talks within cells. Additionally, we report a detailed large-scale biochemical signaling network connecting extra-cellular and cytosolic signals to cytoskeletal dynamics and elements that govern cellular mechanics.

## METHODS

### Building the Cytoskeleton Signaling Network

To build a cytoskeleton signaling network, we extract the data from KEGG pathways^27^ and a variety of related literature. In the following section, we introduce the relationship between cytoskeleton modules and their engagement with upstream biochemical and mechanical cues. All detailed biochemical or mechanical signals and 120 components involved in our model are shown in the **Supplementary Table S1**.

#### Module 1&2: Stress hormones and growth factors control Ca^2+^ homeostasis and GTPases to orchestrate actomyosin signaling

Modules 1 and 2 (Figure 1) describe the relationship between the plasma membrane receptors (PMRs) and actomyosin machinery. Actomyosin elements in the cytoskeleton are involved in cell protrusion, retraction, adhesion, and polarization, and consist of over 100 elements associated with regulating the actin-myosin network (see supplementary for details). PMRs co-operate with GTPases and Ca^+2^ to execute actomyosin structural modifications and functions^37,38^. External signals such as growth factors and stress hormones also use the same machinery to adjust structural elements in the cytoskeleton. Growth factors bind tyrosine kinase receptor (RTK), while adrenergic stress hormones induce G-protein-coupled receptor (GPCR) to coordinate actin polymerization, adhesion and contractility.

To build the cytoskeleton network, we incorporate GTPases upstream of the actin-myosin network such as RhoA, Rac and Cdc42. The findings show that there is complex antagonism between RhoA and Rac. GTPases are induced by various GEF and GAPs as upstream regulators. Such a crosstalk and broad signaling receiver allows fine-tuning of cytoskeletal dynamics required for optimal cell mechanical orchestration and makes GTPases global mediators. RhoA mainly regulates actomyosin contractility and stabilizes actin filaments, promoting highly stable structures such as stress fibers, whereas activation of Rac facilitates lamellipodia formation through actin polymerization.^19,39^ On the other hand, Cdc42 initiates reciprocal signal with Rac, and regulates cell polarity, extension of filipodia and vesicle trafficking ^40,41^.

While GTPases organize cytoskeleton elements in specific manner to generate cell protrusions, Ca^2+^ signaling promotes persistent cell motility. Therefore, we incorporate Ca^2+^ contribution between PMRs and GTPases, or directly on actomyosin network. Ca^2+^ signaling coordinates distinguished local and global pathways. To coordinate cycles in protrusion and retraction, actin polymerization and adhesion formation are regulated by Ca^2+^ release or intake^42^. Additionally, it has been reported that baseline Ca^2+^ concentrations alter Ca^2+^ sensitivity across different signaling pathways and can selectively promote or inhibit actin-myosin dynamics.^43–45^

#### Module 3: Microtubules Signaling

Microtubules (MTs) are dynamic polymers of tubulin within the cytoskeleton and control cell polarization. Apart from the classical role of microtubules in vesicle trafficking, microtubules have recently emerged as key regulators of cell adhesion and migration through their participation in adhesion turnover and cellular signaling. MTs have multiple upstream influencers as shown in supplementary table in microtubule signaling, and in-turn MTs influence actomyosin and focal adhesion interplay through RhoA and TIP+ proteins downstream. Popular known mechanisms facilitating these interactions as upstream signaling include: the activation of Ca^+2^, PI3K, ERK, Cdc42, Rac1 and RhoA signaling by altering MTs between dynamic (polymerizing, depolymerizing and catastrophe) and stable (treadmilling or capped near focal adhesion) states ^46–48^.

#### Module 4: Focal Adhesion Signaling

##### Integrin mechanochemical signaling

Cells sense biochemical and physical signals of the extracellular matrix through adhesive ligands. Integrin-mediated adhesive components interact with specific ECM proteins and sense the rigidity of the substrate to trigger signaling pathways. Integrin activation occurs by outside-in ECM binding or inside-out talin activation which recruits vinculin and F-actins. This process transmits both mechanical (integrin tension dependent dynamics) and chemical signals (focal adhesion kinase and Fyn signaling) from integrins to cytoskeleton, which are incorporated separately in Module1 and Module 4. ECM induced mechanical signals determine the adhesion bound lifetime, whereas FAK phosphorylation provide binding site for Src and other FAK linked proteins ^49^. Phosphorylated FAK promotes F-actin polymerization and regulate contraction by RhoA/ROCK/myosin II complex signaling ^50^.

##### Focal Adhesion Disassembly Signaling

Focal adhesion (FA) disassembly occurs at any stage of adhesion. The disassembly is primarily controlled by tension, endocytosis of integrins and calpain protease activity. Nascent adhesions (NA) generally show a high disassembly rate ^51^. This disassembly is independent from myosin II activity, located in lamella-lamellipodium ^52^. Following this step, integrin clusters in nascent adhesions are either disassembled or mature into focal adhesion complexes. After maturation, mechanical disassembly mostly occurs by MT, FAK or ECM mediated tension (generated via RhoA/ROCK/ppMLC signaling effects on actin-myosin dynamics) on focal adhesion complex in Module 4. High tension on stress fibers ruptures the integrin bond and causes disassembly of FA. In addition to tension dependent dynamics, with chemical signaling, in module 1, Ca^2+^ can bind to Calpain, and cleaves several components of the focal adhesion complex, including talin, paxilin and FAKs ^44^

### BoHyM Model

The base Boolean model comprises of a directed network involving nodes and edges. Every node has its own activator and inhibitor edge as an input; gates between edges, “or” and “and”, provide sufficient or necessary relationship between nodes. In our system, node values show the relative abundance/activity of a component in the network. A node’s state depends on a activation function and decay function which is discussed in detail below. Gates can be “or”, “and” and “or not”. In the depicted network, it is not possible to include an “and not” gate due to the absence of corresponding biological scenarios where proteins have a necessary relationship between both activators and inhibitors.

#### Weight function determines states of the nodes relative to activator or inhibitor density

One of the significant assumptions in Boolean function is that all nodes have equal weightage in turning on succeeding nodes. For examples, if **A** and **B** go to **C** in the network with “or” gate, A and B have will have equal power to make the state of C “1”. However, this is not an exact reflection of biological signals where the networks have ligand affinity and competition between proteins transducing the signals. To account for this biased nature of interactions, we take advantage of a pre-existing model and associated software package, BooleanNet^33^. Across the Boolean models ^53^, **BooleanNet**, a python based software package, provides easiness in modification of piecewise differential equation to adjust relative contribution of preceding signals.

Weight function in our model **Equation {3}** decides strength of activator and inhibitor contribution to succeeding node values. In Boolean models, inhibitors do not show any impact on succeeding node if one of the activators are “on” with all “or” gates. For example, if X_1_^act^ is “on”, it switches the succeeding node’s state independent from the other inhibition edges (Figure 2A). In our model, our weight function reduces the activators’ impact depending on density of inhibitors and activators between the input signals (Figure 2B). Previously Di Cara et al. demonstrated similar approach for weight function in directed network^34^, but in their model Boolean logic has been used to determine the initial states of network nodes and is not applied for general ordinary differential equations, whereas in our model activation function includes weight functions and also logic function with “or” and “and” gates to preserve sufficient and necessity relationship between nodes through simulation. Weight functions range from 0 to 1, where they can reach the maximum value where all activators are 1 without inhibitors, or even with inhibitors, it can asymptotically approach 1 when activator density and values are high enough.

**Figure 2.**
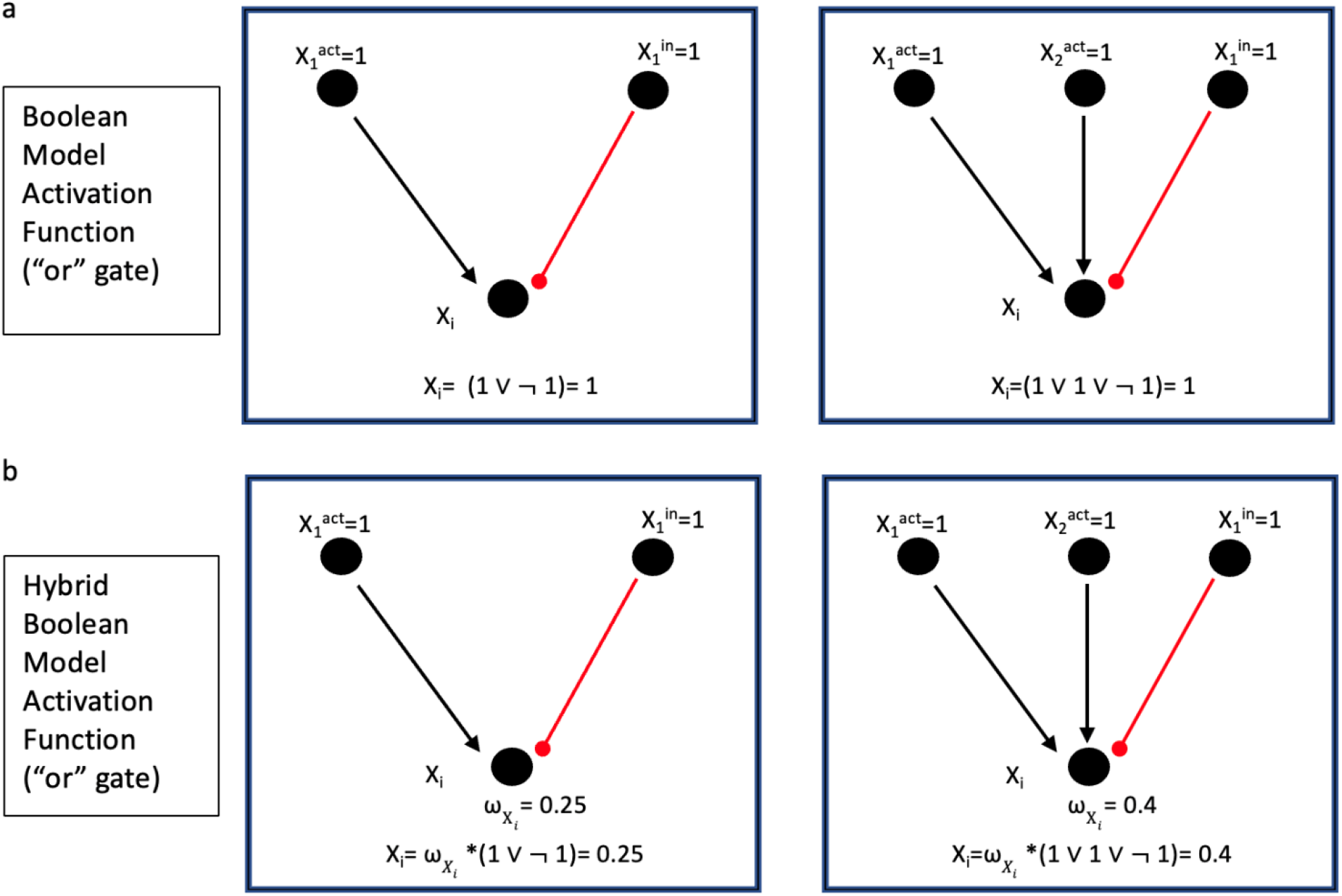
Activation Function of BooleanNet vs. BoHyM. Black arrow: activator; Red pin: inhibitor. (a) In BooleanNet, the activation of a downstream node is computed purely via Boolean logic function. (b) In BoHyM, weight factor, ω_i_, shows total preceding nodes contribution to the activation function, and combined logic function response. X_i_ preceding node values are assigned to maximum level (1.0)

#### Differential equations of BoHyM model

##### 1. Extended Boolean differential equations of general network

To describe the network as a continuous dynamical system, we use the following set of ordinary differential equations:

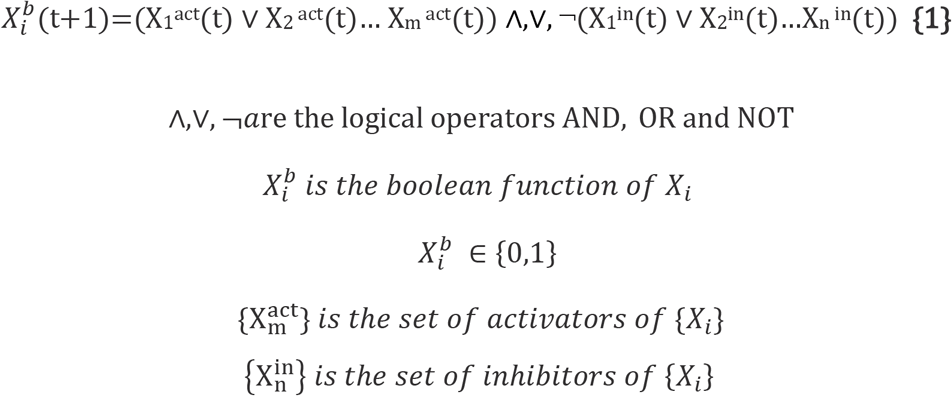

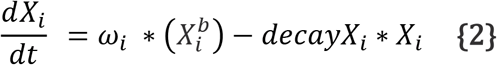

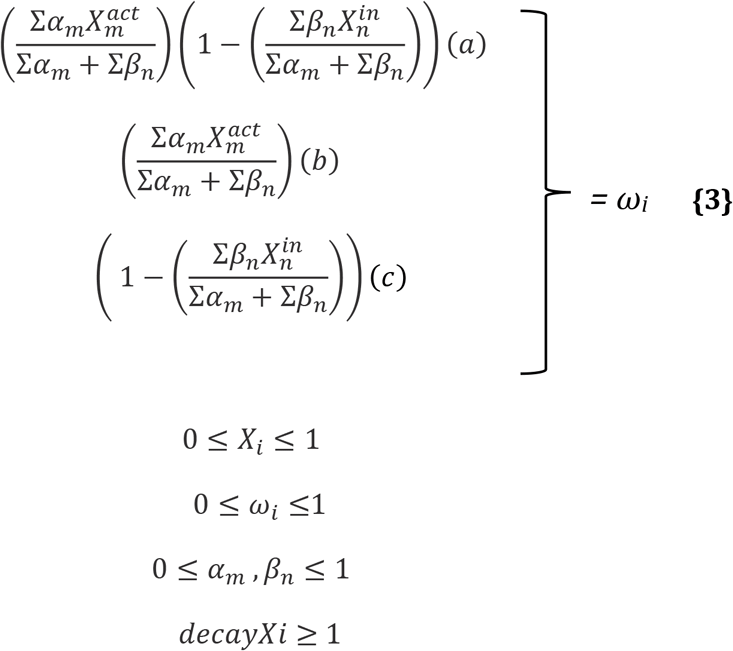

The right-hand side of the differential equations consists of two parts: activation function, and a term for decay, as indicated in BooleanNet. Activation is a multiplication of weight function and Boolean function term which represents the total input to the node. X_i_ is in closed interval [0,1], given that *decayXi is >1 and wi<1.* Subsequently, the second part of the equation is a decay term, which is directly proportional to the level of activation of the node. In all cases, combinations of activators and inhibitors can be also adjusted independently where the adjustment factors are the α and β parameters for activators and inhibitors respectively. The mathematical form of ω_i_ is chosen in the closed interval [0,1], given that 0≤α≤1 and 0≤β≤1. Equations {3a}, {3b} and {3c} show the behavior of ω_i_ when it is controlled subsequently only by combined activators and inhibitors, only activators, or only inhibitors respectively. These Boolean based differential equations describe the states of elements of modules 1 & 2 of the larger cytoskeleton network.

##### 2. Mass-action & Chemo-mechanical Kinetic Models

As described before, Boolean logic cannot capture the all the dynamics of intra-cellular signaling. Therefore, we introduce modular signaling networks in conjunction with the Boolean network.

The first submodule in the cytoskeleton network is MT signaling (module 3), which is integrated into network with mass-action kinetic differential equations. To demonstrate MT dynamics, we split the microtubules states into two groups, shrinking and growing microtubules as dynamic microtubules (dyMT), and stabilized microtubules (stMT). Here total concentration of dyMT and stMT is one, and the mass action kinetics provide dependency between two MTs states. MT states in the submodule are updated through weight functions from module 2, ω_dyMT_ and ω_stMT_ that dictate the state transition rates for the microtubules. The distribution of cytoskeletal microtubules between the two states (dynamic vs stable) is determined using these transition rates as shown in **Equation {4}** & **{5}**. In each time step when analyzing the larger network (modules 1 & 2), weight functions update MT kinetic equations and the submodule (module 3) solves mass-action kinetics, assigning MTs new steady states, which are then passed back to the larger network (module 1) and incorporated as input nodes for downstream Boolean analysis.

Similarly, focal adhesion dynamics are also integrated into model as a second submodule (module 4), where signaling is modeled between integrin, FX and FA. Integrins are activated with talin, vinculin and F-actin to form FX (*reaction ii*); integrin clustering with FX promote FA maturation in (*reaction iii*). In turn FX and FA disassemble into integrin and FX due to tension generated by ECM or ppMLC tension. In focal adhesion kinetic reactions, kβ is the forward reaction rate constant from integrin transformation to focal adhesion complex; ω_ITG_ is dependent on vinculin, talin and actin signals updated from the larger signaling network (**Equation {6})**. kγ is the reaction rate constant for adhesion maturation, and it increases through actin-myosin/ECM generated tension (**Equations {6}, {7} & {8}).** kγ captures the maturation of adhesion by tension with hill functions ^18^. This function reflects the saturation function properties while keeping the constant value between 0-1. It is assumed that the cell starts at a certain minimum level of tension (0.15 value in **Equation {9}**). The other parameters, kδ and kε are disassembly rates in **Equations {10} and {11}**, both are by the function of ECM and Actin-myosin Tension (*X_i_*). Catch-slip dynamics model is used^54^ and the kγ and kε parameters are extracted from Hidalgo-Vasquez study^55^. In catch-slip bond dynamics, as tension increases, bond lifetime has a biphasic trajectory, with lifetime increasing first and subsequently decreasing with applied tension.

In modular networks, values must be scaled to match between the modules. Normalizing in the submodules, kinetic rate constants and weight functions convert the transition values from Boolean scale to mass-action kinetics scale. In both submodules, concentrations of submodule components are normalized by the total concentration. Various factors of X_ECM_ and X_ppMLC_ within **Equations {6}, {7} and {8}** are accordingly selected to match the scaling across the modules. From Boolean to mass-action kinetics ECM are adjusted through the following factors. *X_ECM_A__* = 0.66 * *X_ECM_* + 0.033.

###### Mass-action model

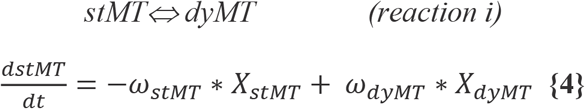

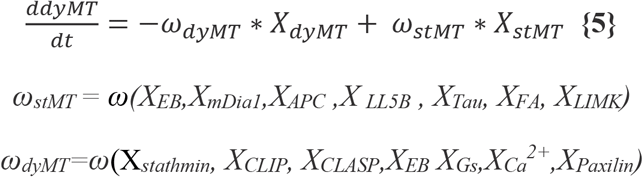

###### Chemo-mechanical model

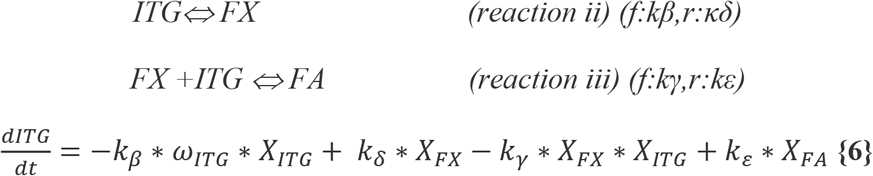

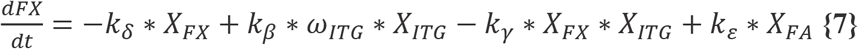

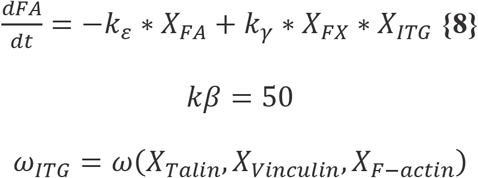

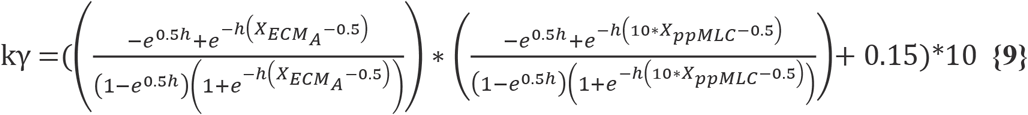

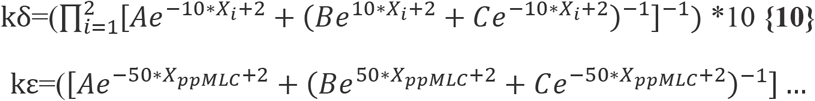

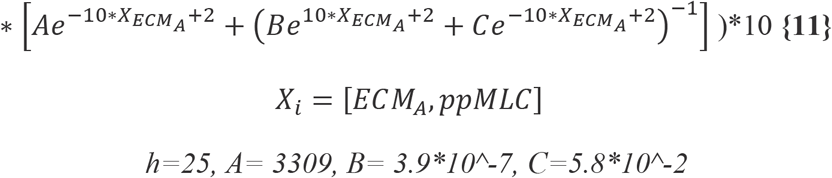

#### Initial States

It is assumed that cells positioned in an extracellular matrix without any cell-cell adhesion consist of active microtubule dynamics and some baseline focal adhesion proteins and focal adhesion complexes. According to the initial conditions, values are assigned to stMT, pip2, integrin, focal adhesion complex, ECM, and certain other nodes in the network that do not have input activators (**Supplementary Figure S1**). Solving the full network with the associated sub-modules provides the values for all nodes which set the reference cell conditions and microenvironment. For the analyses described in this work, the initial values of the input nodes mentioned above are all set at 0.1. These values can be modified by user preferences. The values of all nodes obtained after integrating the BoHyM model for 2000 time points (by which all nodes reach a steady-state) is saved as an initial-states file. Following this step, these steady state node values are always used to initialize the dynamics of signaling for a cell, and only specific nodes are perturbed to simulate different combinations of external signals or changes to intra-cellular expression levels of specific proteins. Perturbations to the node values can be held constant or fluctuated overtime in the model. In the first two applications, perturbation induced in receptor nodes by stress hormones, growth factors and ECM are held constant, assigning values of either 0.1 (low activation) or 1.0 (high activation) to the nodes β_2_AR, ECM, and RTK individually, or in combination with each other. In the last application, states of calcium are temporally modulated to mimic oscillations in intracellular calcium concentrations.

## APPLICATION

### Internal biochemical signals orchestrate the same cytoskeleton elements but reach diverse mechanotypes

Although there are well studied plasma membrane receptors in cellular signaling, it is not quite clear how combining these biochemical cues results in different mechanical properties using similar cytoskeleton elements. Biochemical signals from plasma membrane receptors execute many dynamic changes in the cellular cytoskeleton ^56,57^.Those signals work together in cytoplasm and trigger multiple pathways to change cytoskeleton structure. For instance, GPCRs, Integrins and RTKs share mutual downstream effectors involving PLC, GTPases and Ca^2+^ pathways that regulate cytoskeleton elements (Figure 3A). Tandem impact regarding hormones, growth factor or extracellular matrix level determine the concentration and activity of various cytoskeleton elements and, consequently, mechanical properties of cells^58^.

**Figure 3.**
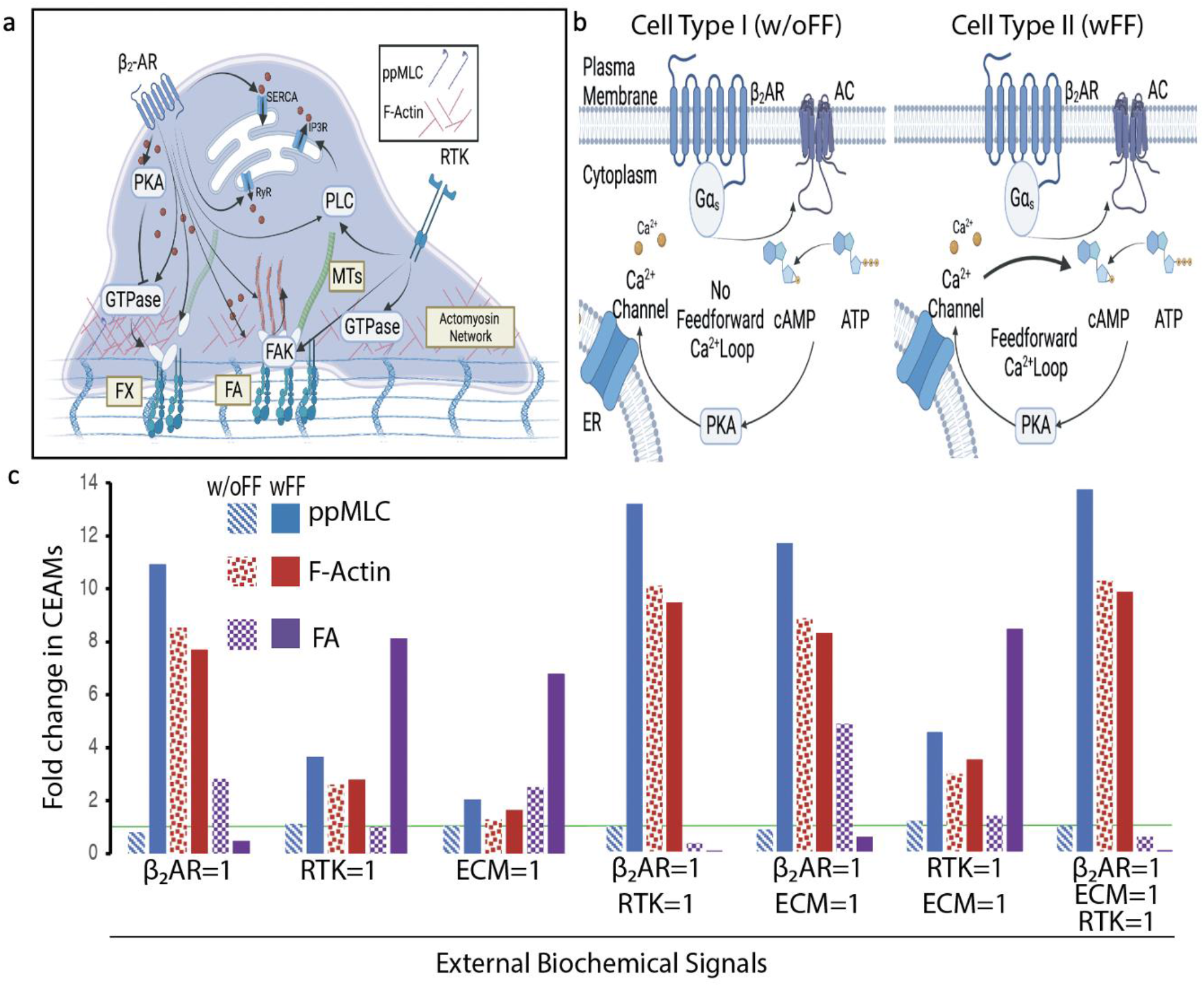
Biochemical signals execute the machinery anchor proteins to modify the mechanical properties of cells. (a) Plasma membrane receptors are rendered here in breast epithelial cells, β_2_-AR sends signal to actomyosin, microtubules, focal adhesion, and calcium channels. Extracellular matrix binds to integrins involved in FX and FA signaling. All receptors use FAK, PLC, GTPase or Ca^2+^ signaling to regulate cytoskeleton elements. (b) Cell type I has no feedforward loop and cell type II has Ca^2+^feedforward loop, which promotes adenylyl cyclase signaling. (c) External biochemical signals modify cytoskeleton components in cells without/with calcium feedforward. Green line represents the initial states. The initial conditions and assumptions for steady states are discussed in methods. Y-axis shows fold change from the initial states in cytoskeleton-ECM anchoring moieties (CEAMs). Left side of Figure C indicates individual β_2_-adrenergic, RTK, ECM perturbations, and the right side is for the combined impact of plasma membrane receptors on cytoskeleton elements. Blue: ppMLC, Red: F-Actin, Purple: Focal Adhesion. Light sheds are for w/o feedforward calcium signaling (cell type I), dark colors are for with feedforward calcium signaling (cell type II)

To investigate how plasma membrane receptor activation changes the cell mechanical properties, we report on the status of three key cytoskeletal-ECM anchoring moieties (CEAMs): filamentous actin (F-actin), di-phosphorylated non-muscle Myosin II light chain (ppMLC) and Focal Adhesion (FA). These components can organize cell behavior by tuning the mechanical properties such as cell stiffness, contractility, and adhesiveness ^59,60^. While several other cytoskeletal proteins also play a significant role in determining cytoskeletal mechanics such as α-actinin, MT end proteins, intermediate filaments, formins and vimentin, we track them within our model, but report only the three CEAMs for the sake of visually representing our results. CEAMs can also be a direct input for some commonly used cellular mechanics and cell migration models ^52,61–65^.

Previous studies indicate that stress hormone signaling leads to various mechanical states in breast cancer cells. Gruet et.al have recently reported that β-adrenergic signaling attenuates the migration in MDA-MB 231, increases migration in MDA-MB-468 and has no change in MCF-7 breast cancer cell lines^66^. Kim et al. show that β-adrenergic agonists promotes migration in MDA-MB 231 cells^67^. Creed et al. also show β-adrenergic ligand increase migration in MDA-MB-231, while Rivero et al. reported the opposite response^68,69^. Therefore, we hypothesize that stress hormones may result in dual mechanical response depending on cell signaling type or cell microenvironment. To test our hypothesis, we investigate slightly varying cell signaling network configurations reported in literature and simulate the downstream effects of β-adrenergic signaling on CEAMs in the presence of different microenvironmental cues. Our model results are employed to further discuss how interventions in certain signaling pathways switch this response and lead to other mechanical responses that are observed in the literature.

Ca^2+^ signaling is known to drive migratory, invasive and proliferative behaviors of tumors ^70–73^. Studies show that less aggressive breast cancer cell lines have lower cytosolic calcium concentrations and an absence of cAMP/Ca^2+^ feedforward loop^38,74,75^. On the other hand, the presence of cAMP/Ca^2+^ feedforward loop is associated with high metastatic potential and invasiveness in cancer cells. Therefore, we define two cell types with slightly different Ca^+2^ signaling structures to observe potential change in cell mechanical properties: Cell type I is without positive feedforward; Cell type II is with positive feedforward (Figure 3B). Both these cell types are initialized as described in the methods section with ECM value set at 0.1, β_2_AR and RTK values set at 0. Using these input values, the initial steady state of the cell types is established by integrating the BoHyM model for 2000 iterations. With these new steady state values as the initial states of the cells, β_2_AR input values are changed from zero to 1 to simulate high-saturated receptors with β-adrenergic agonists. The BoHyM model is simulated for 2000 more iterations with this change to observe the resulting changes in all nodes within the cell signaling network. The steady state results show that β_2_-AR signals downregulate ppMLC and upregulate F-Actin and FA in cell type I, but in contrast increase ppMLC and F-actin, and downregulate FA in cell type II (Figure 3C). BoHyM results indicate that presence of feedforward calcium signaling can reverse the contractility changes observed due to stress hormone signaling in less aggressive breast cancer cell lines which is suggested in previous studies^66,68,76^.

Cells are innately interacting not only with stress hormones but with growth factors and the extracellular matrix in cell micro-environment. It is significant to predict how the combinations of these signaling pathways in conjunction with the β_2_-AR signals influence CEAMs and drive cellular mechanotype responses. Therefore, CEAMs response is analyzed while receptor tyrosine kinase and integrin signaling are activated with different combinations of β_2_-AR signals. Individually high ECM promotes ppMLC only in cell type II, no change in cell type I and increases focal adhesion in both cell types. In between the low and high states of ECM (0.1 – 1.0), it is noticed that ECM modifies FA portion of the CEAMs in a biphasic manner (**Supplementary Figure S2**) along the lines of prior literature ^55,77,78^. On the other hand, RTK signaling increases ppMLC and FA, but the change is more dramatic in cell type II. In addition to individual impact, as a result of receptor crosstalk, it is possible to observe high contractility and weak adhesion when cells are exposed to the stress hormones and growth factor under higher ECM tension. In the absence of stress hormones, both cell types have higher adhesion and lower contractility compared to signals with β_2_AR induction.

### Biased signaling investigation reveals hidden complex dynamics in cytoskeleton network

In complex signaling networks, elements receive inputs from multiple upstream nodes and send outputs to multiple downstream targets. In the results discussed above, it is assumed that all node connections carry equal weights, and the upstream signals influence downstream nodes only according to the activation levels of the upstream nodes and the multiplicity of connections. However, as various signals get distributed and combined along the network, it is not easy to understand the individual contribution of each node to the outcome. To analyze how nodes carry the information from the receptors to various CEAMs, analyzing individual weight functions can be insightful to comprehend the hidden signaling dynamics. Additionally, biases due to the varying interaction affinities between adjacent elements (nodes) can alter the weight of individual connections, beyond just the effect arising from activation levels and multiplicity.

Based on the above arguments, biased distribution between preceding nodes can be classified under two concepts in our model: *In situ* bias and Ligand-based bias. *In situ* bias is the reason for unequal distribution in weight function due to multiplicity of inputs. For instance, there are many GEFs, GAPs and kinases as RhoA activators, and at first glance it is not clear which inputs have a stronger effect between the upstream pathways (Figure 4A). The multiplicity of inputs and outputs into nodes up and downstream of these activators can bias the effects these nodes have on their downstream effectors. Ligand-biased signals occur following modification of α for activators and β for inhibitors values (**Equation 3**). This modification in weight function may mimic biased affinity relationship between ligand receptors and ligands.

**Figure 4.**
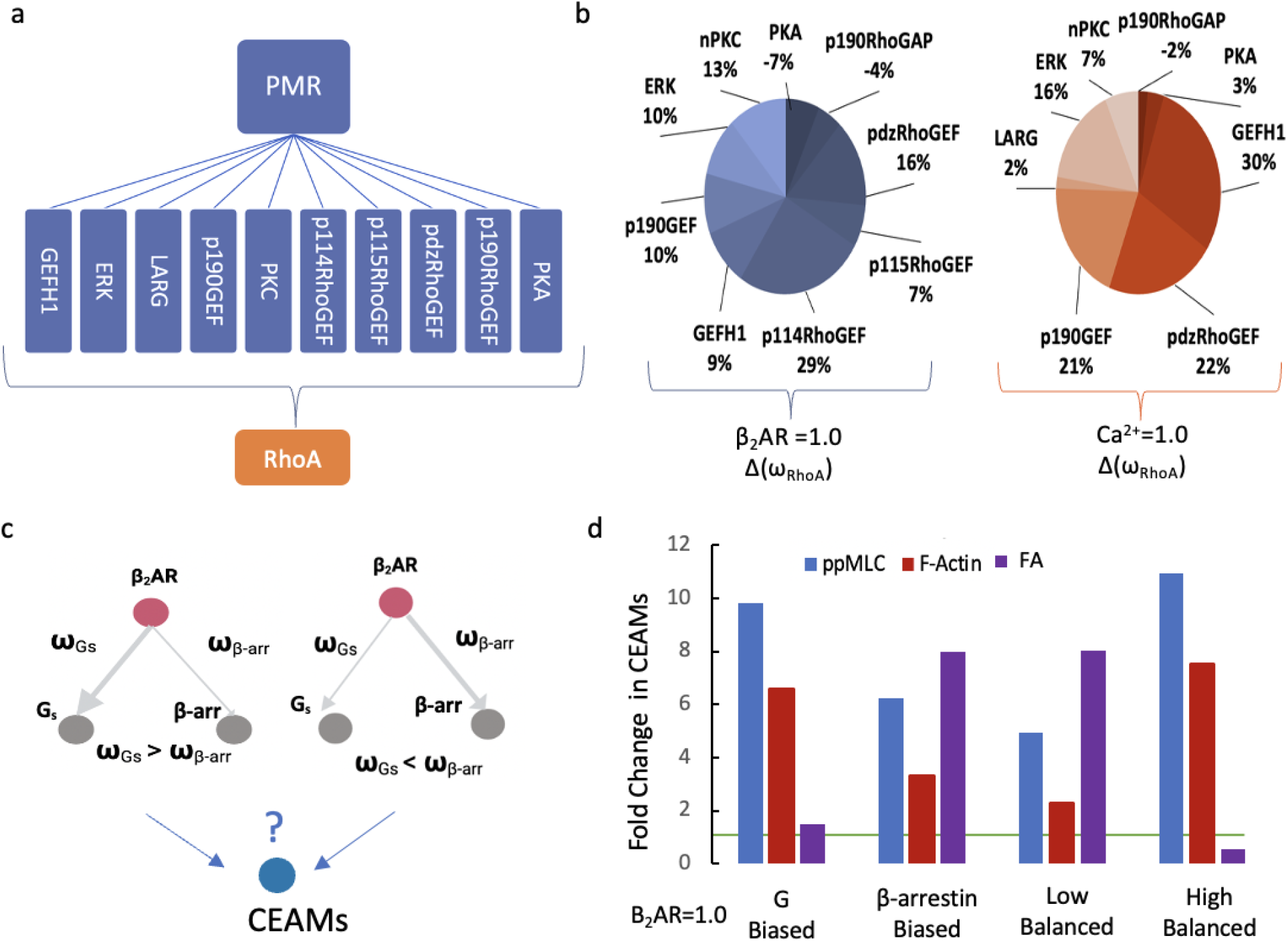
Signaling distribution profile based in situ and ligand biased activation. (a) Plasma membrane receptors (PMR) have multiple inputs into RhoA. (b) shows how β_2_AR and Ca^2+^ contribute to the activation of RhoA at steady states. The values show what percentage of the weight factor driving RhoA signaling is being contributed by the specific preceding nodes indicated in the pie chart. (c) The response of β_2_AR might change due to ligand affinity, it is not clear how this would affect downstream mechanotype dictated by various cytoskeletal elements including CEAMs. (d) In β_2_AR signaling response, the signaling response can be biased by varying affinities in four unique ways - increased G protein affinity, increased β-arrestin affinity, low affinity to both G protein and β-arrestin, and high affinity to both. These differences in ligand affinity can lead to diverse mechanotype profiles as dictated by CEAMs.

To demonstrate application of in situ biased signal analysis, we stimulate β_2_AR and Ca^2+^ signaling as RhoA activators at maximum level (node value set at 1.0). Results from the BoHyM model (Figure 4B) indicate that β_2_-AR modulates RhoA mainly via p114RhoGEF, which receives the signals from Gβγ and β-arrestin (results are shown for Cell Type II). Following p114RhoGEF, pdzRhoGEF, nPKC and p190GEF are the other nodes that have strong impact on RhoA due to network structure. Alternatively, Ca^+2^ drives RhoA activation via GEFH1, pdzRhoGEF and p190GEF pathways (Figure 4B). Ca^2+^ induction promotes GEF-H1 signaling through MT depolymerization, while pdzRhoGEF and p190GEF are mostly initiated by FAK pathways.

It has been shown that β_2_AR has biased signaling depending on ligand availability and distinct response by triggering different pathways such as cAMP/PKA and β-arrestin1/2 ^79^. For mimicking this ligand-biased signaling, α values of β_2_AR in G proteins and β-arrestin weight functions are changed to set a biased relationship between β_2_AR primary messengers. β_2_-AR can activate G-protein signaling and G protein-independent signaling pathways via β-arrestin (Figure 4C). To test the hypothesis that ligand affinity also leads to different response in cytoskeleton network, we increase the activation signal strength for either G-protein or β-arrestin pathways by decreasing the α value in weight function for the complementary pathway (β-arrestin or G-protein respectively) by 10-fold. In figure 4D, using BoHyM, we investigate four cases to show various outcomes of Ligand-biased networks: G-protein biased, β-arrestin1/2 biased, and balanced low/high ligand affinity. Both primary messengers influence cytoskeleton mechanical properties individually in a similar manner, just at different amplitudes (results are shown for Cell Type II). Collectively however, we observe that G proteins increase contractility strongly and focal adhesions weakly, whereas β-arrestin signaling impacts for contractility and focal adhesions strongly. When G protein and β-arrestin signaling are balanced, having a low ligand affinity moderately increases contractility and strongly increases adhesion. In contrast, a higher ligand affinity strongly increases the contractility but decreases focal adhesions.

### BoHyM can simulate oscillatory mechanical responses based on transient signals

Averaged over intermediate timescales (on the order of minutes), concentrations of specific proteins and small molecule effectors seem to hold steady within the cell. However, at shorter timescales (on the order of microseconds to seconds), levels of proteins and ions can oscillate significantly. These transient signals are necessary for a cell’s repetitive movements and continuous controlling mechanisms against external and internal cues ^80–82^. Using transient signals, cells orchestrate their movements and adjust their cytoskeleton properties driving protrusion and retraction dynamics driving mechanosensing and migration ^54,83–87^.

As one example of these transient signals, Ca^2+^ pulses have a major role in orchestrating cell cytoskeleton dynamics and provide quick mechanical responses in short intervals of time. Pulses of Ca^2+^ coordinate actin assembly for stepwise cell extension^88^. Calcium elevations increase the residency of FAK at FAs, thereby leading to increased activation of its effectors involved in FA disassembly^89^. Moreover, calcium waves regulate RhoA/ROCK contractility through microtubules depolymerization^90^. It has been shown that calcium signaling can adapt MLCK transient signaling and local cycles of lamellipodia retraction and adhesion in migrating cells^91^. Thus, cells modulate protrusion dynamics and the rate of cell migration in response to the amplitude and frequency of calcium signaling via a variety of interacting pathways. Understanding the coupling between Ca^+2^ signaling and cytoskeletal dynamics is important in predicting mechanosensory and migratory response of cells to transient Ca^+2^ signaling in specific biochemical and mechanical environments.

To investigate the relationship between the calcium pulses and protrusion chemo-mechanical dynamics, we analyze the elements of cytoskeleton upon being affected by transient Ca^2+^ signals. Ca^2+^ signals are induced sinusoidally in the model. Our findings indicate that there is a synchronized oscillation between myosin phosphorylation and calcium signaling through MLCK, PAK and ROCK pathways. Because of their dependence on acto-myosin contractility, FA adhesions also oscillate, however FA formation and dissolution are shifted in relation to calcium dynamics (Figure 5B). PAK has dual pathways, whereby it increases contractility through Cdc42 and Rac upstream signaling, while also inhibiting MLCK following calcium increase. We believe this duality leads to secondary small waves in focal adhesion dynamics observed in Figure 5B. As a simplified extension of focal adhesion dynamics, we assume that cell speed during cell crawling is dictated by the rate of focal adhesion turnover ^82^. Based on this, we can predict cell fluctuations at the front of a migrating cell as a function of Ca^+2^ dynamics. We observe interesting bimodal fluctuations in cell front speeds in response to each calcium pulse, which may correlate to certain experimental observation reported in literature^92,93^. While we do not claim to recreate or explain any specific experimental observations, our results display the ability of the BoHyM model to describe experimentally observed temporal cytoskeletal dynamics as a function of transient signaling in cells.

**Figure 5.**
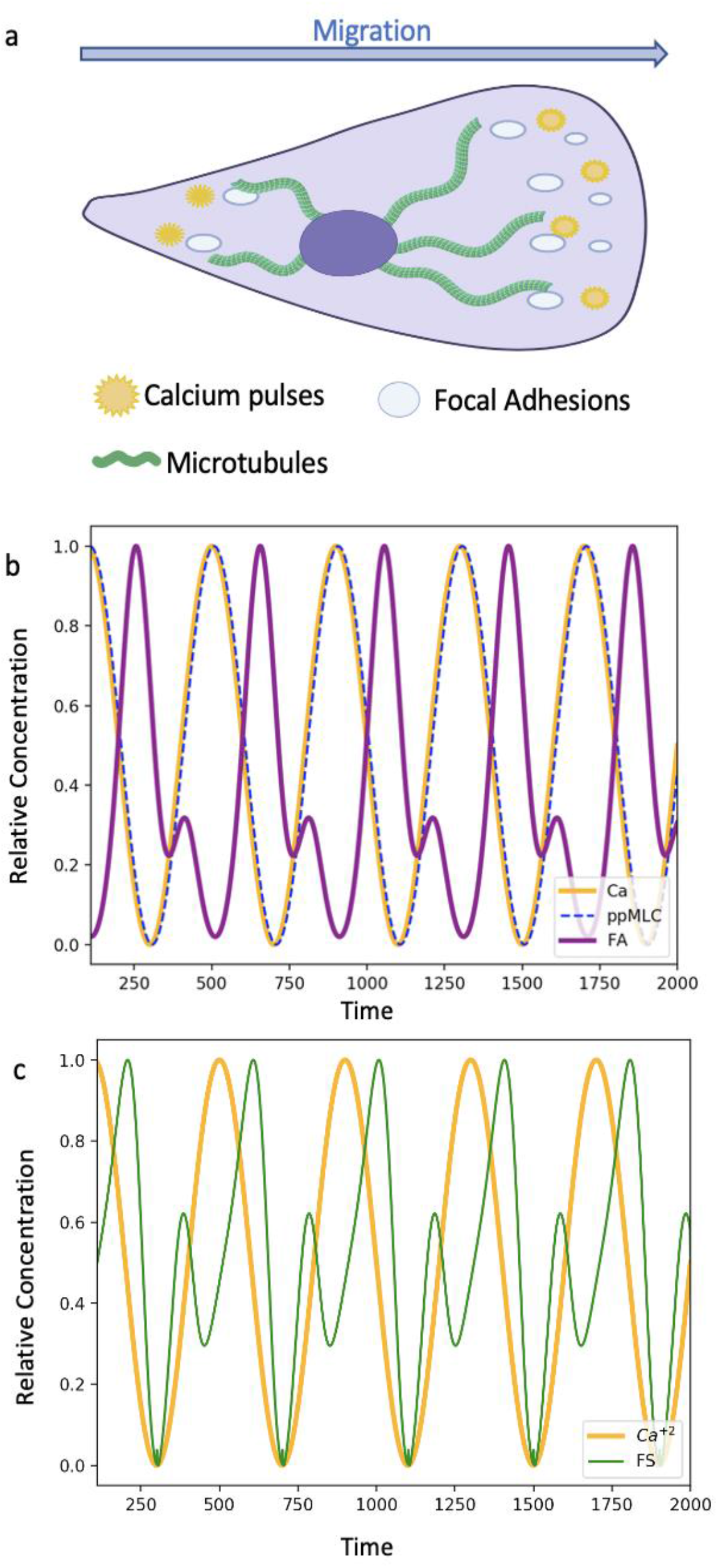
Ca^2+^ oscillations control protrusion dynamics through cytoskeleton elements. (a) This figure shows that calcium concentration has gradient in front and rear of cells. Calcium pulses promote retraction and propulsion in the leading protrusions (b) Orange line: Ca^2+^, blue dash line: ppMLC, purple line: FA, this figure demonstrates how intra-cellular calcium fluctuation is reflected through myosin phosphorylation and focal adhesion concentration changes over time. (c) Orange line: Ca^2+^, Green line: Front speed. Migrating cell’s Front speed is calculated as a function of the focal adhesion turnover rate (dFA/dt) and its relationship to Ca^2+^ cycles is shown.

## DISCUSSION

Biochemical signaling affecting cytoskeleton dynamics is vast and complicated. Kinetic models require precise rate parameters. Boolean models simplify the network with logic functions in possible modules. Adding a few kinetic mass-action module overcomes Boolean limitations with relatively fewer and measurable kinetic parameters. Using our model, we show how external signals including growth factors, β-adrenergic stress hormones and ECM stiffness, and internal protein expression levels influence various cytoskeletal elements in phenotypically different cell types.

BoHyM provides a modular framework for changing the structure and weights rapidly to match experimental conditions. Specifically, nodes and connections can be optimized to replicate specific experimental/clinical outcomes. In addition, the model allows forward integration of biochemical signal to mechanical models. Key nodes that are input for those models are contractility, ECM stiffness, focal adhesions, traction forces and focal adhesion turnover rates. The model also allows backward integrations of mechanical signals from the environment, neighboring cells etc. to study how these translate into cellular signaling through ECM, integrin, cadherin & tension nodes.

In this work we have built the architecture of the cytoskeletal signaling network to contain network connections and logical combinations based on what we were able to confirm through prior literature (see Supplementary Table S1). However, a more comprehensive combinatorial analysis for network connections and interaction types can be performed to understand in-depth how biological signaling may regulate the entire spectrum of cell mechanical behaviors. Due to the large number of elements accounted for in the current signaling network, and a lack of complementary biological validation for predicted outcomes at the moment, we believe this task to be out of the scope for this manuscript, but something that we and others can explore in the future.

One limitation of our model is that it does not include spatial resolution in cell signaling and temporal resolution at multiple time scales. Spatial models are necessary to define delays/spatial hinderances in transmitting signals^94^. Spatial modules with temporal delays between signals transmitted and received between modules seem to be an easily deployable and viable candidate to incorporate spatial dynamics within the current BoHyM architecture. Another limitation in signaling models is integrating signaling events over multiple timescales. For example, a lot of the upstream signaling affecting cytoskeletal elements and specific downstream signaling based on the current state of these cytoskeletal elements can drive transcription factor activation and altered gene signaling. However, these signaling events occur at relatively longer timescales which are not being explored in the current version of our model. Extension of BoHyM model with spatial models and signaling at different time scales will provide an even more comprehensive platform to capture complex interactions in intracellular signaling.

Overall, we have established a first of its kind Boolean-Hybrid-Modular (BoHyM) model to study biochemical regulation of cytoskeletal mechanics that incorporates multiple cell-environment interactions as well as extensive intra-cellular signaling. The model incorporates 120 individual signaling elements, 279 connections between these elements, MT dynamics and chemo-mechanosensitive FA formation and dissolution. Additional modules for transcription level signaling, cell membrane mechanics, cytoskeletal-nuclear mechanics and so on can be added for additional realism or to answer specific questions.

There is a growing need for modular models providing cell signaling while considering chemo-mechanical interactions. It is important to remember that mechanistic and biochemical signaling components are inseparable. Additionally, new biophysical approaches are identifying complex regulations and interaction between the chemical signals a cell receives, the biochemical state of the cell, the mechanical signals a cell receives, the mechanical state of the cell and how all of these combine to generate a complex chemo-mechanical response landscape for cell behavior in various biological processes. What we have provided here is an important tool integrating and analyzing these complex interactions and predicting cell mechanotype based on cell phenotype and the microenvironment.

## CONCLUSION

We have developed a semi-quantitative-computational model to analyze the intra-cellular signaling network and its outcome in the presence of multiple external signals including growth factors, hormones, and extracellular matrix. We use this model to analyze the cell response phase space to external stimuli and identify the key internal elements of the network that drive specific outcomes. The model is built upon Boolean approach to network modeling. This allows us to analyze the network behavior without the need to estimate all the various interaction rates between different cellular components. However, since our approach is limited in its ability to predict network dynamics and temporal evolution of the cell state, we introduce dynamical aspects using mass-action kinetics for signal transmission rates within the network as well as chemo-mechanical models on kinetic data. Combining these approaches, we provide a unique computational model to connect cells biochemical signaling profile to mechanical properties which are key to understanding cellular force generation and migration in response to those mechanical and chemical cues.

## ACKNOWLEDGEMENT

Thanks to Dr. Parag Katira, CompActMatter Lab and Dr. Stephanie I. Fraley, Fraley Lab for their support and feedback. Dr. Katira acknowledges funding from the NSF CMMI (Award #1905390) for this research. Dr. Stephanie I. Fraley acknowledges funding from NSF CAREER (1651855 to S.I.F). BioRender.com and yEd-graph editor were used to render images and networks graphs.

## AUTHOR CONTRIBUTION

E.T.K., S.I.F, and P.K. conceived of the project and designed *the in-silico* experiments. All computational modeling was performed by E.T.K., A.T and J. G. helped analyze model outputs and test specific scenarios. The manuscript was written by E.T.K., A.T., S.I.F, and P.K. with input from the other authors.

## CODE AVALIBILITY

The python code used to simulate the cytoskeleton signaling and the model is available via GitHub (https://github.com/etkarabay/BoHyM-boolean-hybrid-modular-model). The code is divided into two different sections. Initial values generator can be found under the section labeled initial states file. Cell type I/II simulator are available under the example file.

