## Supplementary Information for "Predicting phenotype to mechanotype relationships in cells based on intracellular signaling network"

### Supplementary Figures

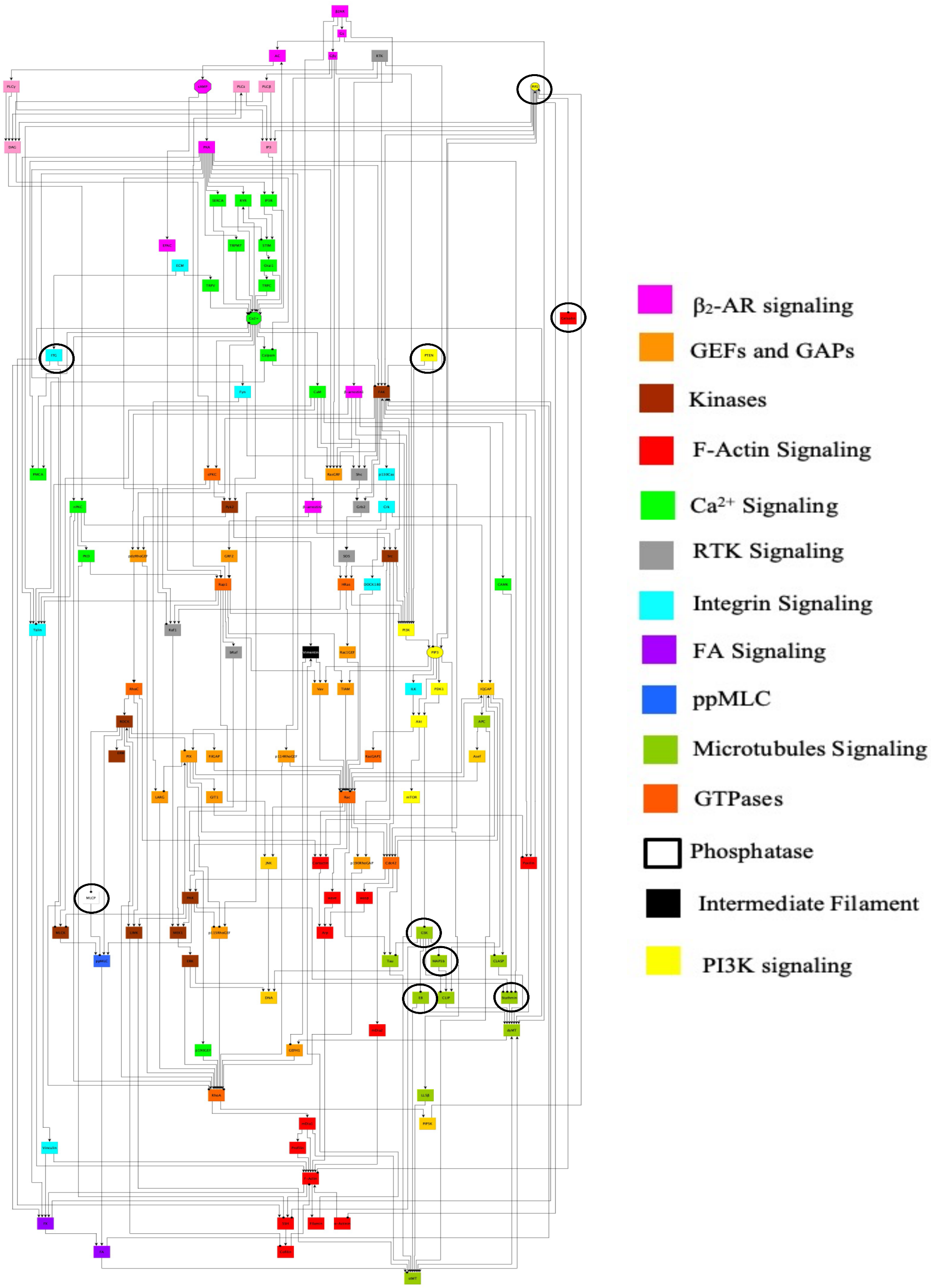


**<u>Figure S1. All cytoskeletons network.</u>** Colors represent different signaling modules. The shapes other than rectangular shows the non-protein components such as $Ca^{2+}$ and cAMP. Black circled nodes show the components initiated at 0.1 as mentioned methods. yEd-graph editor is used to render the interactive network.

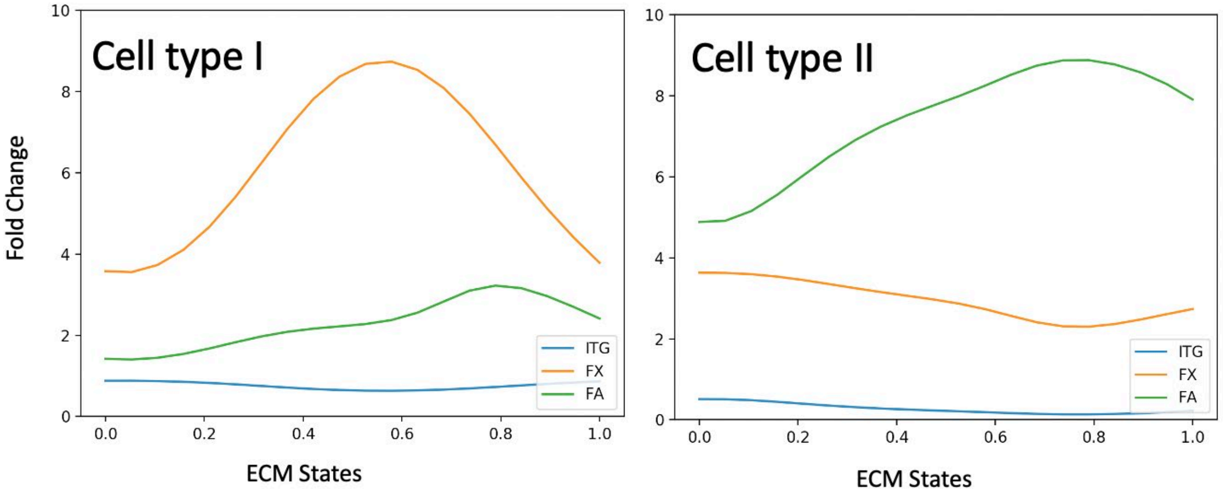


**Figure S2. Focal adhesion components have biphasic response regarding ECM states.**

Depending on the ECM states integrins, focal adhesion complexes (FX) and focal adhesions (FA) show biphasic dynamics. Blue Line: Integrin, Orange Line: Focal adhesion complexes Green Line: Focal Adhesion

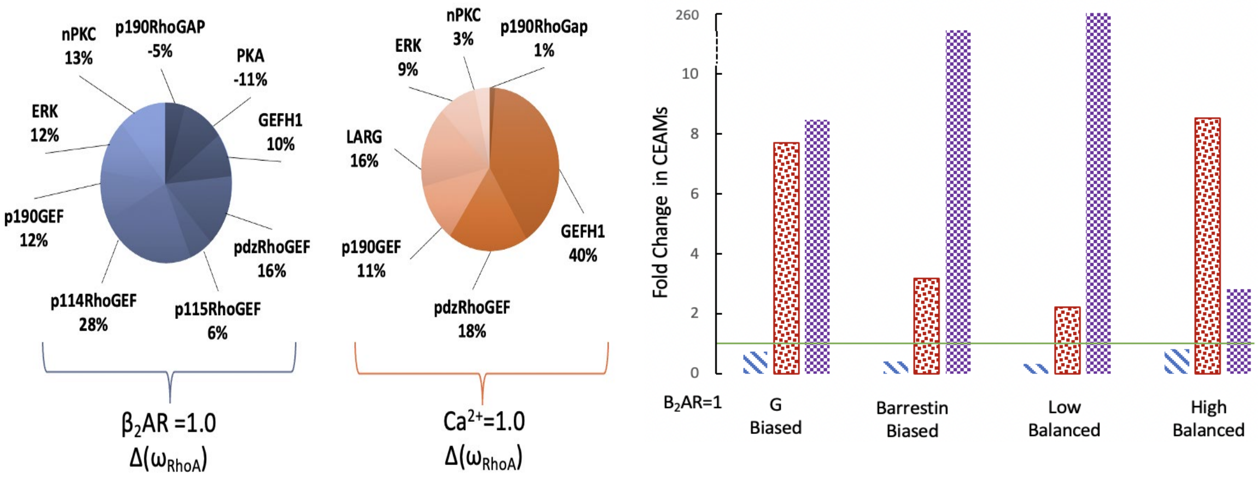


**Figure S3. Cell Type I Signaling distribution profile based in situ and ligand biased activation.**

(a) It shows how $\beta_2$AR and $Ca^{2+}$ contribute to the activation of RhoA at steady states in Cell Type I. The values show what percentage of the weight factor driving RhoA signaling is being contributed by the specific preceding nodes indicated in the pie chart. (b) In $\beta_2$AR signaling response, the signaling response can be biased by varying affinities in four unique ways - increased G protein affinity, increased β-arrestin affinity, low affinity to both G protein and β-arrestin, and high affinity to both. These differences in ligand affinity can lead to diverse mechanotype profiles as dictated by CEAMs.

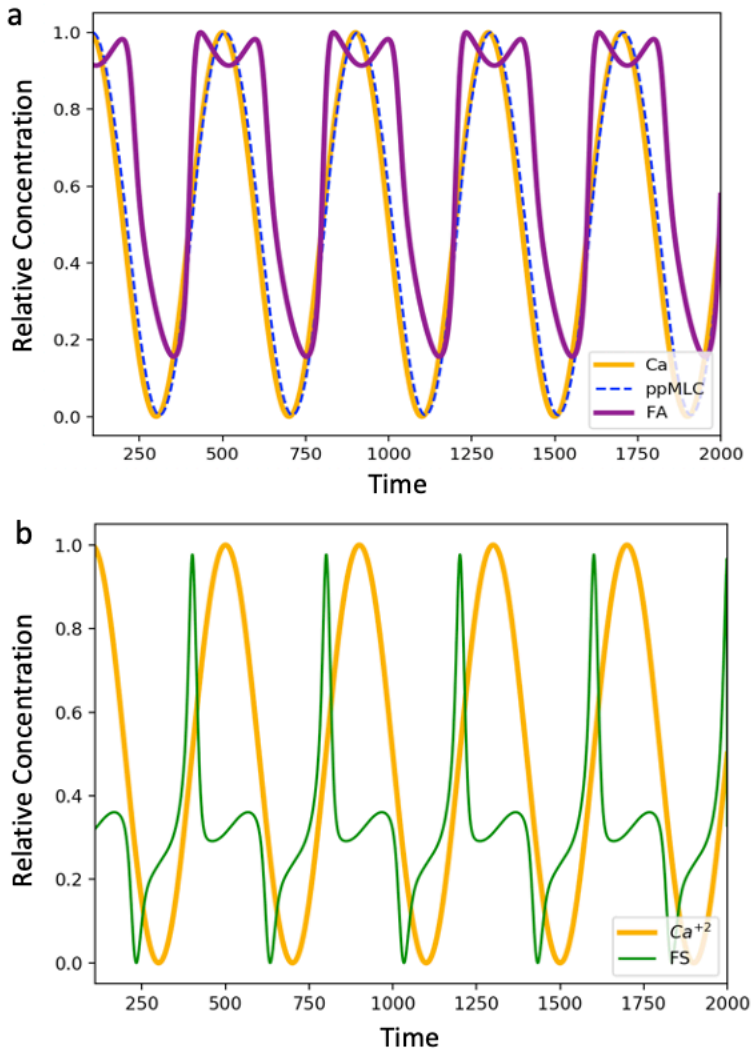


**Figure S4. Cell Type II oscillatory mechanical responses based on transient signals.**

Orange line: $Ca^{2+}$, Green line: Front speed. Migrating cell's front speed is calculated as a function of the focal adhesion turnover rate (dFA/dt) and its relationship to $Ca^{2+}$ cycles is shown in Cell Type II

**Supplementary Tables**

**Table S1. Cytoskeleton Directed Network**

**Calcium Signaling Network**

| Signaling Pathway | References |
|---|---|
| PIP2 → IP3 | [1] |
| PLCg → IP3 | Calcium Signaling Pathway [2] |
| PLCe → IP3 | Calcium Signaling Pathway [2] |
| PLCb → IP3 | Calcium Signaling Pathway[2] |
| PTEN --\| PIP3 | Calcium Signaling Pathway[2] |
| PIP2 → PIP3 | Calcium Signaling Pathway[2] |
| PI3K → PIP3 | Calcium Signaling Pathway[2] |
| DAG → nPKC | Calcium Signaling Pathway[2] |
| Ca2+ → cPKC | Calcium Signaling Pathway[2] |
| PLCg → DAG | Calcium Signaling Pathway[2] |
| PLCb → DAG | Calcium Signaling Pathway[2] |
| PLCe → DAG | Calcium Signaling Pathway[2] |
| PIP2 → DAG | [1] |
| IP3R → $Ca^{2+}$ | Calcium Signaling Pathway [2] |
| $Ca^{2+}$ → Calpain | [3] |
| PKA --\| Calpain | [4] |
| PKA → TRPM7 | [5] |
| PKA → IP3R | [6] |
| IP3 → IP3R | Calcium Signaling Pathway [2] |
| STIM → Orai1 | Calcium Signaling Pathway [2] |
| Orai1 → $Ca^{2+}$ | Calcium Signaling Pathway [2] |
| RyR → STIM | Calcium Signaling Pathway [2] |
| IP3R → STIM | Calcium Signaling Pathway [2] |
| $Ca^{2+}$ → RyR | Calcium Signaling Pathway [2] |
| $Ca^{2+}$ → CaM | Calcium Signaling Pathway [2] |
| $Ca^{2+}$ → Pyk2 | Calcium Signaling Pathway [2] |
| CaM → CAMK | Calcium Signaling Pathway [2] |
| SERCA --\| $Ca^{2+}$ | Calcium Signaling Pathway [2] |
| RyR → $Ca^{2+}$ | Calcium Signaling Pathway [2] |
| PMCA --\| $Ca^{2+}$ | Calcium Signaling Pathway [2] |
| SERCA --\|STIM | Calcium Signaling Pathway [2] |

| | |
|---|---|
| CaM → RasGRF | RasGRF Signaling Pathway[7] |
| ECM → TRPV | [8] |
| Orai1 → TRPC | [9] |
| CaM → PI3K | [10] |
| TRPV → $Ca^{2+}$ | [8] |
| TRPC → $Ca^{2+}$ | [9] |
| TRPM7 → $Ca^{2+}$ | [5] |
| Barrestin1 --\| RasGRF | [11] |
| FAK → RasGRF | Focal Adhesion Signaling[2] |
| CaM →PMCA | [12] |
| PKA → SERCA | Calcium Signaling Pathway [2] |
| Rap1 → PLCe | cAMP Signaling Pathway[2] |

**Microtubule Network**

| | |
|---|---|
| Stathmin → dyMT | [13] |
| CLIP → dyMT | [14] |
| CLASP → dyMT | [14] |
| EB →dyMT | [14] |
| Gs → dyMT | [15] |
| $Ca^{2+}$ → dyMT | [16] |
| Paxilin → dyMT | [17] |
| PKA --\| Stathmin | [18] |
| PAK --\| Stathmin | [19] |
| ERK --\| Stathmin | [20] |
| CAMK --\| Stathmin | [21] |
| nPKC → IQGAP | [22] |
| Barrestin1 → IQGAP | [22] |
| IQGAP → CLIP | [23] |
| Map1b → CLIP | [24] |
| IQGAP → CLASP | [25] |
| Vimentin --\| GEFH1 | [26] |
| dyMT → GEFH1 | [26] |
| GSK --\| Tau | Alzheimer Disease Signaling Pathway[2] |
| Rac → Tau | [27] |
| Cdc42 → Tau | [28] |
| LL5β → stMT | [29] |
| CLASP → stMT | [29] |
| EB → stMT | [30] |
| mDia1 → stMT | [31] |
| APC → stMT | [31] |
| Tau → stMT | [32] |
| FA → stMT | [33] |
| LIMK --\| stMT | [34] |
| CLIP → stMT | [35] |
| PIP3 → LL5β | [36] |
| GSK --\| EB | [37] |
| GSK --\| MAP1B | [38] |

**Focal Adhesion Network**

| | |
|---|---|
| ECM → ITG | Focal Adhesion Pathway[2] |
| ITG → Src | Focal Adhesion Pathway[2] |
| ITG →Fyn | Focal Adhesion Pathway[2] |
| Src → FAK | Focal Adhesion Pathway[2] |
| FAK → Src | Focal Adhesion Pathway[2] |
| cPKC → FAK2/Pyk2 | Focal Adhesion Pathway[2] |
| Src → p190RhoGAP | Focal Adhesion Pathway[2] |
| FAK → p190GEF | [39] |
| PIP5K → PIP2 | Focal Adhesion Pathway[2] |
| ROCK → ppMLC | Focal Adhesion Pathway[2] |
| ROCK --\| MLCP | Focal Adhesion Pathway[2] |
| MLCP --\|ppMLC | Focal Adhesion Pathway[2] |
| RhoA → mDia1 | Focal Adhesion Pathway[2] |
| FAK → Paxilin | Focal Adhesion Pathway[2] |
| FAK →PI3K | Focal Adhesion Pathway[2] |
| FAK → p130Cas | Focal Adhesion Pathway[2] |
| PTEN --\| FAK | Focal Adhesion Pathway[2] |
| PIP2 → Vinculin | Focal Adhesion Pathway[2] |
| Calpain --\| Talin | Focal Adhesion Pathway[2] |
| Crk → Paxilin | Focal Adhesion Pathway[2] |
| p130Cas →Crk | Focal Adhesion Pathway[2] |
| PIP3 →ILK | Focal Adhesion Pathway[2] |
| ILK → Akt | Focal Adhesion Pathway[2] |
| PIP3 → PDK1 | Focal Adhesion Pathway[2] |
| PDK1 →Akt | Focal Adhesion Pathway[2] |
| PIP3 → Vav | Focal Adhesion Pathway[2] |
| Vav → Rac | Focal Adhesion Pathway[2] |
| Rac → PAK | Focal Adhesion Pathway[2] |
| Crk → GRF2 | Focal Adhesion Pathway[2] |
| Crk → DOCK180 | Focal Adhesion Pathway[2] |
| DOCK180 → Rac | Focal Adhesion Pathway[2] |
| GRF2 → Rap1 | Focal Adhesion Pathway[2] |
| Rap1 → B-Raf | Focal Adhesion Pathway[2] |
| B-Raf → MEK1 | Focal Adhesion Pathway[2] |
| PAK → MEK1 | Focal Adhesion Pathway[2] |
| MEK1 → ERK | Focal Adhesion Pathway[2] |

| | |
|---|---|
| Rap1 → JNK | Focal Adhesion Pathway[2] |
| PAK --\| MLCK | Focal Adhesion Pathway[2] |
| RhoA → ROCK | Focal Adhesion Pathway[2] |
| RhoA → PIP5K | Focal Adhesion Pathway[2] |
| Cdc42 → PAK | Focal Adhesion Pathway[2] |
| Rac → JNK | Focal Adhesion Pathway[2] |
| PIP3 → Akt | Focal Adhesion Pathway[2] |
| Pyk2 → Src | Focal Adhesion Pathway[2] |
| Cdc42 → mDia2 | [40] |
| Src → PI3K | [1,241] |
| PKA → Talin | [42] |
| cPKC → Talin | [42] |
| PIP2 → Talin | [42] |
| Rap1 → Talin | [42] |
| Talin → Vinculin | Focal Adhesion Pathway[2] |
| Paxilin → Vinculin | Focal Adhesion Pathway[2] |
| Vimentin --\| GEFH1 | [26] |
| Fyn → LARG | [43] |
| PAK → p115RhoGEF | [44] |
| PKA --\| RhoA | cAMP Signaling Pathway[2] |
| GEFH1 → RhoA | [45] |
| pdzRhoGEF → RhoA | [46] |
| p115RhoGEF → RhoA | [47] |
| p114RhoGEF → RhoA | [48] |
| p190RhoGAP → RhoA | Focal Adhesion Pathway[2] |
| p190GEF → RhoA | [49] |
| LARG → RhoA | [43] |
| pdzRhoGEF → RhoC | [50] |
| F-Actin → FX | [51] |
| Vinculin →FX | [51] |
| Talin → FX | [51] |
| RhoC → ROCK | [52] |
| PKA → FAK | [53] |
| PIP2 → FAK | [54] |
| FX/FA → FAK | [54] |

**RTK Signaling**

| | |
|---|---|
| RTK → Shc | Focal Adhesion Pathway[2] |
| Shc → Grb2 | Focal Adhesion Pathway[2] |
| FAK → Grb2 | Focal Adhesion Pathway[2] |
| FAK → Shc | Focal Adhesion Pathway[2] |
| RTK → FAK | Focal Adhesion Pathway[2] |
| Grb2 → SOS | Focal Adhesion Pathway[2] |
| SOS → HRas | Focal Adhesion Pathway[2] |
| HRas → Raf1 | Focal Adhesion Pathway[2] |
| Raf1 → MEK1 | Focal Adhesion Pathway[2] |
| RTK → PI3K | Focal Adhesion Pathway[2] |
| H-Ras → PI3K | Focal Adhesion Pathway[2] |
| Fyn → Shc | Focal Adhesion Pathway[2] |
| ERK → DNA | Focal Adhesion Pathway[2] |
| mTOR → DNA | Focal Adhesion Pathway[2] |
| JNK → DNA | Focal Adhesion Pathway[2] |
| Akt → mTOR | Focal Adhesion Pathway[2] |
| RTK → PLCγ | Calcium Signaling Pathway[2] |
| Src → HRas | [55] |
| Src → PI3K | [56] |
| cPKC → Raf1 | [57] |
| nPKC → Vimentin | [58] |
| Filamin → Vimentin | [59] |
| nPKC → PKD | Rap Signaling Pathway[2] |
| ERK → RhoA | [60] |
| nPKC → RhoA | [61] |
| p114RhoGEF → Rac | [48] |
| PAK → ppMLC | [62] |

**Actin Cytoskeleton Signaling**

| | |
|---|---|
| LARG → RhoA | Regulation of Actin Cytoskeleton[2] |
| HRas → Rac1GEF | Regulation of Actin Cytoskeleton[2] |
| Rac1GEF → Rac | Regulation of Actin Cytoskeleton[2] |
| Vav → Rac | Regulation of Actin Cytoskeleton[2] |
| Tiam1 → Rac | Regulation of Actin Cytoskeleton[2] |
| Cdc42 → Wasp | Regulation of Actin Cytoskeleton[2] |
| Cdc42 → IQGAP | Regulation of Actin Cytoskeleton[2] |
| Rac → IQGAP | Regulation of Actin Cytoskeleton[2] |
| PIX --\| LARG | Regulation of Actin Cytoskeleton[2] |
| PAK → PIX | Regulation of Actin Cytoskeleton[2] |
| PIX → GIT1 | Regulation of Actin Cytoskeleton[2] |
| PIX → Cdc42 | Regulation of Actin Cytoskeleton[2] |
| PIX → Rac | Regulation of Actin Cytoskeleton[2] |
| mDia1 → Profilin | Regulation of Actin Cytoskeleton[2] |
| PIP2 → Vinculin | Regulation of Actin Cytoskeleton[2] |
| PIP2 --\| Actinin | Regulation of Actin Cytoskeleton[2] |
| GIT1 --\| Paxilin | Regulation of Actin Cytoskeleton[2] |
| ROCK → ERM | Regulation of Actin Cytoskeleton[2] |
| ROCK → LIMK | Regulation of Actin Cytoskeleton[2] |
| LIMK --\| Cofilin | Regulation of Actin Cytoskeleton[2] |
| SSH → Cofilin | Regulation of Actin Cytoskeleton[2] |
| PIP2 --\| Gelsolin | Regulation of Actin Cytoskeleton[2] |
| Rac → Wave | Regulation of Actin Cytoskeleton[2] |
| Asef → Rac | Regulation of Actin Cytoskeleton[2] |
| APC → Asef | Regulation of Actin Cytoskeleton[2] |
| Wave → Arp | Regulation of Actin Cytoskeleton[2] |
| Wasp → Arp | Regulation of Actin Cytoskeleton[2] |
| PIP3 → TIAM | Regulation of Actin Cytoskeleton[2] |
| Src → Cortactin | [63] |
| F-Actin → Actinin | [63] |
| Cortactin → Arp | [63] |
| Vimentin → Vav | [64] |
| PAK → LIMK | Regulation of Actin Cytoskeleton[2] |
| Rac → Cortactin | [63] |
| Actin → Filamin | [65] |
| PKD --\| SSH | [66] |

| | |
|---|---|
| ROCK --\| PIX | [67] |
| PKA → PIX | [68] |
| ROCK → FilGAP | [69] |
| Profilin → F-Actin | Regulation of Actin Cytoskeleton[2] |
| Arp → F-Actin | Regulation of Actin Cytoskeleton[2] |
| Rac1 --> p190RhoGAP | [70] |
| Cofilin --\| F-Actin | Regulation of Actin Cytoskeleton[2] |
| Gelsolin --\| F-Actin | Regulation of Actin Cytoskeleton[2] |
| IQGAP → F-Actin | [23] |
| mDia1 → F-Actin | Regulation of Actin Cytoskeleton[2] |
| mDia2 → F-Actin | Regulation of Actin Cytoskeleton[2] |
| Rac → Cdc42 | Regulation of Actin Cytoskeleton[2] |
| Src →Cdc42 | [71] |
| FilGAP --\| Rac | [69] |
| PKA → PIX | [68] |
| GSK --\| SSH | [72] |
| $Ca^{2+}$ → SSH | [73] |
| F-Actin → SSH | [74] |
| cPKC → Cdc42 | [75] |
| RasGRF →HRas | Ras Signaling Pathway |

**$\beta_2$AR Signaling**

| | |
|---|---|
| Ca → AC | cAMP Signaling Pathway[2] |
| PKA → PMCA | cAMP Signaling Pathway[2] |
| PMCA --\| Ca2+ | cAMP Signaling Pathway[2] |
| PKA → RyR | cAMP Signaling Pathway[2] |
| PKA --\| RhoA | cAMP Signaling Pathway[2] |
| PKA --\| Raf1 | cAMP Signaling Pathway[2] |
| BAR2 → Gs | cAMP Signaling Pathway[2] |
| Gs → AC | cAMP Signaling Pathway[2] |
| AC → cAMP | cAMP Signaling Pathway[2] |
| cAMP → PKA | cAMP Signaling Pathway[2] |
| cAMP → EPAC | cAMP Signaling Pathway[2] |
| EPAC → Rap1 | cAMP Signaling Pathway[2] |
| Rap1 → PI3K | cAMP Signaling Pathway[2] |
| Rap1 → TIAM | cAMP Signaling Pathway[2] |
| Rap1 → Vav | cAMP Signaling Pathway[2] |
| Barrestin2 → Src | [76] |
| BAR2 → Barrestin1/2 | [77] |
| BAR2 → Gbg | [78] |
| Barrestin2 → p115RhoGEF | [79] |
| p114RhoGEF → Rac | [48] |
| Gbg → p114RhoGEF | [48] |
| PAK → ppMLC | [80] |
| Barrestin1 → pdzRhoGEF | [81] |
| Pyk2 → p190GEF | [39] |
| FAK → p190RhoGEF | [39] |
| pyk2 → pdzRhoGEF | [46] |
| FAK → pdzRhoGEF | [82] |
| Gbg → PI3K | Pathways in Cancer[2] |
| Barrestin1 → PI3K | [83] |
| Rap1 → Raf1 | Rap1 Signaling Pathway[2] |
| PKD → Rap1 | Rap1 Signaling Pathway[2] |
| Gbg → PLCe | [84] |
| PKA → RasGRF | Ras Signaling Pathway[2] |
| Gbg → RasGRF | Ras Signaling Pathway[2] |

**Table S2. Components Names and Symbols**

| Symbols | Name | Also known as |
|---|---|---|
| **RacGAP1** | Rac GTPase activating protein 1 | |
| **Barrestin1** | Beta Arrestin 1 | |
| **Barrestin2** | Beta Arrestin 2 | |
| **RhoC** | Ras Homolog Family Member C | |
| **nPKC** | Novel Protein Kinase C ( δ,ε,η,θ) | |
| **cPKC** | Conventional Protein Kinase C(α ,βI,βII γ) | |
| **Cortactin** | Cortactin | |
| **Vinculin** | Vinculin | |
| **RasGRF** | Ras protein specific guanine nucleotide releasing factor 1 | |
| **TRPM7** | Transient Receptor Potential Cation Channel Subfamily M Member 7 | |
| **PMCA** | Plasma Membrane Ca2+ ATPase | |
| **Filamin** | Filamin | |
| **Vimentin** | Vimentin | |
| **PKD** | polycystin-1 | |
| **FS** | Front Speed | |
| **Pyk2** | Protein Tyrosine Kinase 2 | PTK2 |
| **Calpain** | Ca2+-activated neutral cysteine proteases | |
| **p115RhoGEF** | Rho Guanine Nucleotide Exchange Factor 1 | ARHGEF1 |
| **p114RhoGEF** | Rho Guanine Nucleotide Exchange Factor 18 | ARHGEF18 |
| **pdzRhoGEF** | PDZ Rho Guanine Nucleotide Exchange Factor | |
| **Gbg** | G beta-gamma complex | |
| **EPAC** | Rap Guanine Nucleotide Exchange Factor 3 | RAPGEF2 |
| **STIM** | Stromal Interaction Molecule | |
| **Orai1** | Calcium Release-activated calcium channel protein 1 | |
| **RyR** | Ryanodine Receptors | |
| **SERCA** | Sarco-/Endoplasmic Reticulum Ca2+ ATPase | |
| **Fyn** | Photo-oncogene tyrosine-protein kinase Fyn | |
| **ITG** | Integrins | |
| **LARG** | Leukemia-associated Rho Guanine Nucleotide Exchange Factor | ARHGEF12 |
| **Asef** | Rho Guanine Nucleotide Exchange Factor 4 | ARHGEF4 |
| **ERM** | Ezrin, Radixin, Moesin | ARHGEF28 |
| **p190GEF** | p190 Rho Guanine Nucleotide Exchange Factor | RGNEF (mouse) |

| | | |
|---|---|---|
| **FilGAP** | Filamin A Binding RhoGTPase Activating Protein | |
| **SSH** | Protein Phosphatase Slingshot | |
| **IQGAP** | IQ motif-containing GTPase-activating Protein | |
| **p190RhoGa p** | p190 Rho Family GTPase-activating Protein | GRLF1 |
| **GIT1** | ARF GTPase-activating Protein | ARFGAP1 |
| **mDia2** | Mouse Diaphanous 2 | |
| **PLCe** | Phospholipase C-epsilon | |
| **PLCg** | Phospholipase C-gamma | |
| **FA** | Focal Adhesion | |
| **FX** | Focal Adhesion Complex | |
| **APC** | Adenomatous Polyposis Coli | |
| **GEFH1** | Rho Guanine Nucleotide Exchange Factor 2 | ARHGEF28 |
| **ECM** | Extracellular Matrix | |
| **CLASP** | CLIP-associating proteins | |
| **dyMT** | Dynamic Microtubules | |
| **Tau** | microtubule stabilizer protein | |
| **stMT** | Stable Microtubules | |
| **LL5B** | Phosphatidylinositol (3,4,5) Trisphosphate Sensor proteins | |
| **EB** | End-binding Protein | |
| **MAP1b** | Microtubule Associated Protein 1B | |
| **CLIP** | Class II-associated invariant chain peptide | |
| **PTEN** | Phosphatase and Tensin Homolog | |
| **Rac1GEF** | Rac1 GTPase Guanine Nucleotide Exchange Factor | |
| **Gelsolin** | Gelsolin | |
| **Profilin** | Profilin | |
| **PIX** | p21-actciated Protein Kinase Exchange Factor | ARHGEF7 |
| **Stathmin** | Stathmin | |
| **PDK1** | Pyruvate Dehydrogenase Kinase1 | |
| **Paxilin** | Paxilin | |
| **Talin** | Talin | |
| **Actinin** | Alpha-actinin-1 | |
| **ILK** | Integrin-linked protein kinase | |
| **ppMLC** | di-phosphorylated non-muscle myosin light chain | |
| **Src** | proto-oncogene c-Src | |
| **MLCK** | Myosin Light Chain Kinase | |
| **cAMP** | cyclic adenosine monophosphate | |
| **p130Cas** | p130 CRK-associated Substrate | |
| **GRF2** | General Regulatory Factor 2 | |

| | | |
|---|---|---|
| **IP3** | Inositol Triphosphate | |
| **MLCP** | Myosin Light Chain Phosphatase | |
| **TIAM** | T-cell lymphoma invasion and metastasis-inducing protein 1 | |
| **F-Actin** | Actin Filaments | |
| **bRaf** | b-RAF proto-oncogene | |
| **Cofilin** | Cofilin | |
| **RhoA** | Ras Homolog Family Member A | |
| **PAK** | p21 activated kinases | |
| **Vav** | Rho Guanine Nucleotide Exchange Factor | |
| **GSK** | Glycogen synthase kinase 3 | |
| **ERK** | Extracellular signal regulated kinase | MAPK |
| **DAG** | Diglyceride | |
| **PIP5K** | Phosphatidylinositol 4-phosphate 5-kinase | |
| **Rap1** | Saccharomyces cerevisiae repressor-activator protein 1 | |
| **PIP2** | Phosphatidylinositol 4.5-bisphosphate | |
| **MEK1** | Mitogen -activated protein kinase 1 | MAP2K |
| **RTK** | Receptor Tyrosine Kinase | |
| **Wave** | WASP-family verprolin-homologous protein | |
| **FAK** | focal adhesion kinase | |
| **SOS** | Son of Sevenless | |
| **Grb2** | Growth Factor Receptor Bound Protein 2 | |
| **mTOR** | Mammalian Target of Rapamycin | |
| **ROCK** | Rho-associated protein kinase | |
| **AC** | adenylate cyclase | |
| **DNA** | Signals change cell proliferation or cell cycle | |
| **PIP3** | Phosphatidylinositol 3,4.5-triphosphate | |
| **PI3K** | Phosphoinositide 3-kinases | |
| **HRas** | RasGTPase | |
| **mDia1** | Mouse Diaphanous 2, Formin | |
| **Akt** | Protein Kinase B | |
| **DOCK180** | Dedicator of cytokinesis | DOCK1 |
| **Shc** | Shc-transforming protein 1 | |
| **PKA** | Protein Kinase A | |
| **Crk** | Adapter Molecule Crk | |
| **Raf1** | Raf-1 photo-oncogene, serine/threonine kinase | |
| **β2AR** | β2- Adrenergic Receptor | |
| **Gs** | Gs alpha subunit | |
| **Rac** | Rho Family GTPases | |
| **LIMK** | LIM Domain Kinase 1 | |

| | | |
|---|---|---|
| **Ca** | Calcium | |
| **Wasp** | Wiskott-Aldrich Syndrome Protein | |
| **JNK** | c-Jun N-terminal Kinases | |
| **Arp** | Actin Related Protein 2/3 | ARP2/3 |
| **Cdc42** | Cell Division Cycle 42 | |
| **CaM** | Calmodulin | |
| **CaMK** | Calmodulin Dependent Protein Kinase | |
| **TRPV** | Transient Receptor Potential Cation Channel V | |
| **TRPC** | Transient Receptor Potential Cation Channel C | |
| **PLCb** | Phospholipase C-beta | |
| **IP3R** | Inositol Trisphosphate Receptor | |
